# Assessing Forearm Exertion in Manual Tasks with Surface EMG: A Comparative Analysis of Through-Forearm vs. Muscle Specific EMG placements

**DOI:** 10.1101/2024.12.10.627589

**Authors:** Xuelong Fan, Johan Rydgård, Liyun Yang, Peter J. Johansson

## Abstract

**Background:** Hand-intensive work is associated with musculoskeletal disorders (MSDs) such as carpal tunnel syndrome, tendinitis, and nerve entrapment. Estimating internal load on tissues like tendons and muscles in the hand and forearm is essential but challenging. Consequently, external force or self-reported exertion are often used as proxies for internal load assessment. Surface electromyography (sEMG) provides a promising method for continuous and objective monitoring of force. Through-forearm sEMG – placing one electrode on the flexors and one on the extensors – has been suggested as a pragmatic approach for capturing physical load under different wrist and forearm postures. However, further validation is needed to determine its accuracy in predicting exertion and force across diverse tasks, as well as to understand what specific aspects of muscle activity it captures. This study aimed to compare through-forearm sEMG with two common muscle-specific placements, i.e., the forearm extensor and flexor, in estimating force and perceived exertion during hand-intensive tasks and to explore the correlations of sEMG signals between these three sEMG placements.

**Methods:** Sixteen participants performed four simulated tasks, i.e. gripping, thumb-pressing, screwing, and key-pinching, at five exertion levels following a modified OMNI-Resistance Exercise Scale (OMNI-RES). The sEMG signals from the through-forearm, extensor, and flexor placements were recorded alongside exerted force. Polynomial mixed-effects models were used to estimate self-rated exertion and exerted force from muscle activity. Pearson correlation and linear mixed-effects models were used to explore correlations of sEMG signals between these three sEMG placements.

**Results:** All sEMG placements predicted self-rated exertion and exerted force with strong model fit (R^2^ > 0.95 in most tasks) and high precision (residual SD < 1 for self-rated exertion, and < 5% for force in most tasks). However, moderate inter-subject and inter-task variability was observed. Through-forearm sEMG consistently outperformed both extensor and flexor placements across tasks by a small margin. The sEMG signals obtained from the three sEMG placements were highly correlated, and through-forearm sEMG could be expressed as a linear combination of extensor and flexor signals.

**Discussion:** The findings suggest that exertion estimation with a single-channel sEMG is feasible, although individual calibrations and additional task-specific information may be necessary for real-world applications. Through-forearm sEMG placement marginally outperformed extensor and flexor placements, which could be due to its integration of muscle activities from both muscle groups.

**Conclusion:** The through-forearm sEMG placement provides slightly better performance estimating exertion than muscle-specific placements in the assessed manual tasks, possibly due to its integration of information from multiple muscles. Further research is needed to explore methods for individual calibration and ways to obtain task-specific information to enhance applicability in broader real-world settings.

## 1 Introduction

Musculoskeletal disorders (MSDs) of the hand and arm are a prevalent issue, negatively impacting wellbeing (Renée Govaerts et al., 2021), life expectancy (Ritsuno et al., 2021), and productivity (Epstein et al., 2018), while also incurring great economic costs (Power et al., 2022). Workers involved in hand-intensive tasks are particularly susceptible to developing MSDs, such as carpal tunnel syndrome (Palmer et al., 2006), tendinitis (Van Rijn et al., 2008), and nerve entrapments (Arvidsson et al., 2003; Van Rijn et al., 2008).

To mitigate the risk of MSDs, regular risk assessments are essential to identify and address potentially hazardous work activities, in accordance with health and safety legislation and established guidelines (Council of the European Union, 1989). Accurate assessment of physical workload is a key factor in preventing MSDs (Van Der Beek et al., 2017). Various risk assessment tools (Bonfiglioli et al., 2013; Jorgensen et al., 2024; Steven Moore & Garg, 1995) use scale-based methods to estimate force exposure and temporal information, which are subsequently integrated into overall MSD risk assessments. While these tools provide valuable insights, they often rely on self-reports or third-party observations, which may disrupt ongoing work and limit the number and length of tasks that can be evaluated. Additionally, scale-based tools are commonly known to be tedious and resource-consuming, which is not suitable for long-term and continuous monitoring.

To overcome these challenges, sensor-based assessment tools, such as inertial measurement units (IMUs) and surface electromyography (sEMG), have been introduced to reduce observer biases and workload, enable efficient continuous monitoring, and save time and resources (Balogh et al., 2009; Hansson et al., 2006; Nordander et al., 2004). Kinematic outcomes measured by IMUs can effectively reflect an individual’s exposure to awkward postures and hazardous fast or repetitive movements, making them useful for assessing the risks of MSDs (Arvidsson et al., 2021). However, kinematic measurements lack direct information on the actual loads in the musculoskeletal system, making it difficult to distinguish between forceful and non-forceful tasks, which limits their applicability in many occupational settings. Directly estimating internal load on tissues like tendons and muscles in the hand and forearm would be ideal but challenging. Consequently, external force or self-reported exertion are often used as proxies for internal load assessment. As a result, there remains a need for reliable, efficient, and unobtrusive tools to directly monitor and record physical load during work tasks.

Surface electromyography (sEMG) offers potential to address this gap by estimating physical load continuously over extended periods with minimal interference with work tasks (Fan et al., 2022; Hansson et al., 2009, 2010). sEMG signals have been shown to correlate with hand exertion force (Barański et al., 2024; Bardizbanian et al., 2020; Mao, Fang, et al., 2023), providing critical missing information on the force for risk assessment. However, various factors, including wrist and forearm postures (Takala & Toivonen, 2013), types of gripping (Greig & Wells, 2008), and muscle selection (Forman et al., 2021) contribute to variability in the relationship between force demands and measured muscle activity. While combining multiple muscle signals can improve the accuracy of predictions (Martinez et al., 2020; Tepe & Demir, 2022), it also increases the complexity of the system and application. The limited surface area of the forearm further restricts the number of possible sEMG electrode placements, posing additional challenges. Although integrated multi-channel sEMG systems, such as one-dimensional ring arrays, have demonstrated sufficient accuracy in predicting hand gestures (Tepe & Demir, 2022), their high cost and limited commercialization make them impractical for widespread use in clinical or occupational settings.

Takala and Toivonen (2013) proposed a pragmatic alternative by measuring surface sEMG through the forearm (through-forearm) using a single bipolar channel, with one electrode on the extensor and one on the flexor muscle groups. This approach demonstrated more consistent exertion load estimations across various wrist and forearm postures compared to muscle-specific placements, such as those targeting only the extensor or flexor. The through-forearm approach appears to offer a viable alternative for assessing the general physical load on the forearm and hand in occupational environments.

However, research on through-forearm sEMG is limited. The existing study by Takala and Toivonen (2013) did not examine the performance of the through-forearm placement during other common work-related tasks, such as twisting and pinching, nor did it evaluate the ability of through-forearm sEMG to predict exerted force. Therefore, further investigation is necessary to examine the performance of through-forearm sEMG for assessing the exerted force in a wider range of hand-intensive tasks.

This study aims to:

1. Investigate whether the through-forearm sEMG placement provides better and more consistent estimates of exertion across different tasks compared to the muscle-specific sEMG placements on the extensor and flexor muscles. The measures of exertion include:

a. Self-reported perceived exertion
b. Exerted force
2. Explore the correlations between the through-forearm sEMG placement and the extensor and flexor placements to understand the muscle activity captured by the through-forearm sEMG.

The two-fold aims are also illustrated in Figure 1.

**Figure 1.**
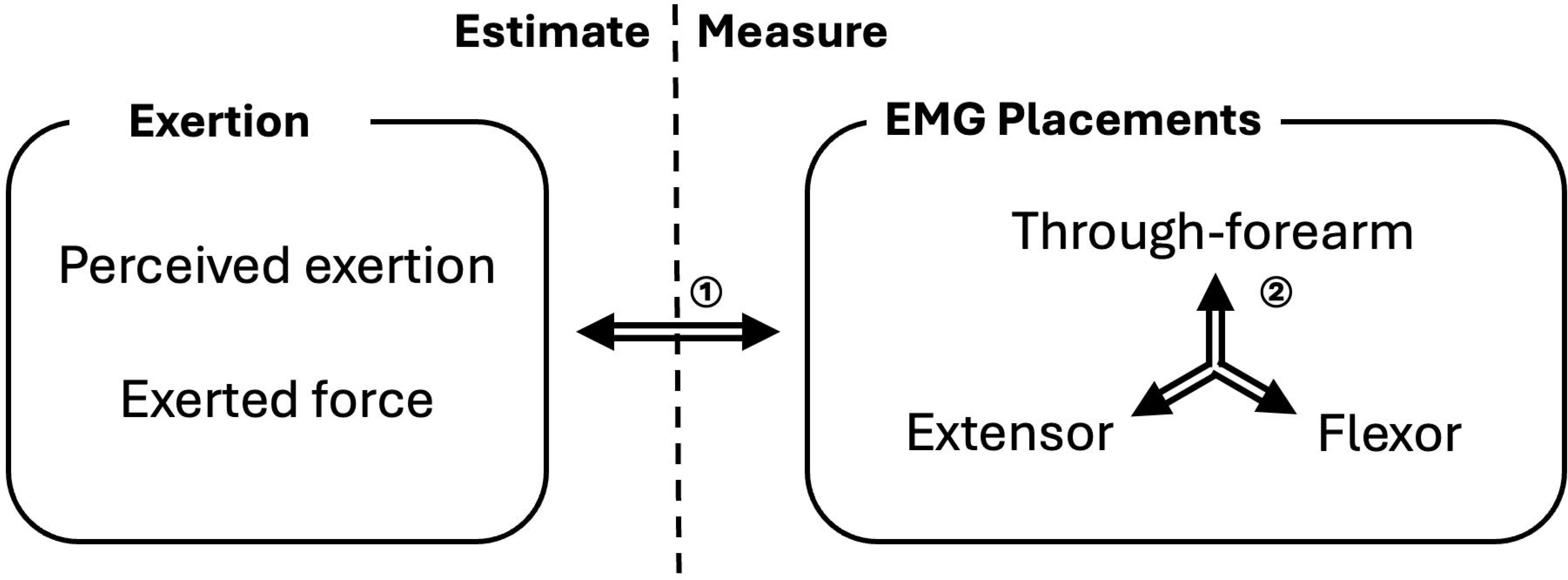
The scheme of the study. Each multidirectional arrow corresponds to one aim of the study.

## 2 Methods and materials

### 2.1 Participants

Sixteen healthy, female (56%) and male (44%) participants (mean ± SD: age, 41 ± 10 years; height, 173 ± 9 cm; BMI, 24 ± 3 kg/m^2^; wrist circumference, 166 ± 13 mm), either right- (75%) or left-handed (25%), from a Swedish research institute participated the study voluntarily (Table 1). All participants were informed of the studies and have given their consent. The study was approved under ethical approval by The Ethical Review Board in Gothenburg (Ref. no. 2022-02531-01) and funded by AFA Insurance Ref.no. 200070.

**Table 1.** Description of the participants (N=16).

| Characteristics | Value |
| --- | --- |
| Age, mean (SD), years | 41 (10) |
| Female, N (%) | 9 (56) |
| Height, mean (SD), cm | 173 (9) |
| Weight, mean (SD), kg | 73 (11) |
| BMI, mean (SD), kg/m<sup>2</sup> | 24 (3) |
| Wrist circumference, mean (SD), mm | 166 (13) |
| Right-handed, N (%) | 12 (75) |

### 2.2 Experiment settings

The study included four predefined simulated manual tasks: gripping, thumb-pressing, screwing, and key-pinching. The experiment consisted of two phases: calibration and task.

The **calibration phase** included one resting period and four maximal voluntary contraction (MVC) periods (one for each task). The **resting period** took place at the beginning of the experiment. During the resting period, participants first stood upright with their eyes facing forward and arms hanging naturally at their sides for 15 seconds. They were then seated in a chair with back support, with the dominant arm placed on an armrest at a 90° elbow angle for 15 seconds. The height of the armrest was adjusted to ensure that the shoulder remained relaxed. This resting period was used to determine the noise level for the sEMG measurements, which could be obtained from either the standing or sitting position (See Section 2.5). The four **MVC periods** were conducted at the start of each task. During these periods, participants were instructed to conduct the corresponding task by gradually increasing their exertion to the maximum, under verbal encouragement from a researcher, and holding this maximal voluntary contraction for 3 seconds before relaxing (at least 30 seconds). The four MVC periods were used to normalize the exerted force for the four tasks. Noticeably, the MVC period for gripping was also used to normalize EMG signals (See 2.5 Signal processing below).

In the **task phase**, participants performed the four predefined tasks. The order of tasks was fixed to minimize the number of participant groups required, and at least 2 minutes of rest was provided between the tasks. Before each task, participants positioned their dominant hand on a corresponding exertion meter in the appropriate posture and practiced the task until they felt ready. They then rested their hand in position without exerting force for 10 seconds to prepare for the task session. Following the MVC period for each task and a resting period of at least 30 seconds, participants were asked to increase their force to three predetermined perceived exertion levels, corresponding to the scores of 2, 4, and 8 on an OMNI-RES scale that was modified to better illustrate hand exertion (Figure 2). A 10 on the modified OMNI-RES is defined as the maximal exerted force and a zero is defined as 5% of the maximal exerted force or below. The rest numbers, e.g., 1, 2, 3, etc., represent 10%, 20%, 30% of the maximal exerted force and so on. The participants were holding the level for 5 seconds before relaxing. A minimum resting period of 20 seconds was provided between each exertion. To reduce the number of required participant groups, the exertion levels were presented in a hierarchically randomized order, with each participant assigned to one of four sequences: 2-4-8, 4-2-8, 8-2-4, or 8-4-2.

**Figure 2.**
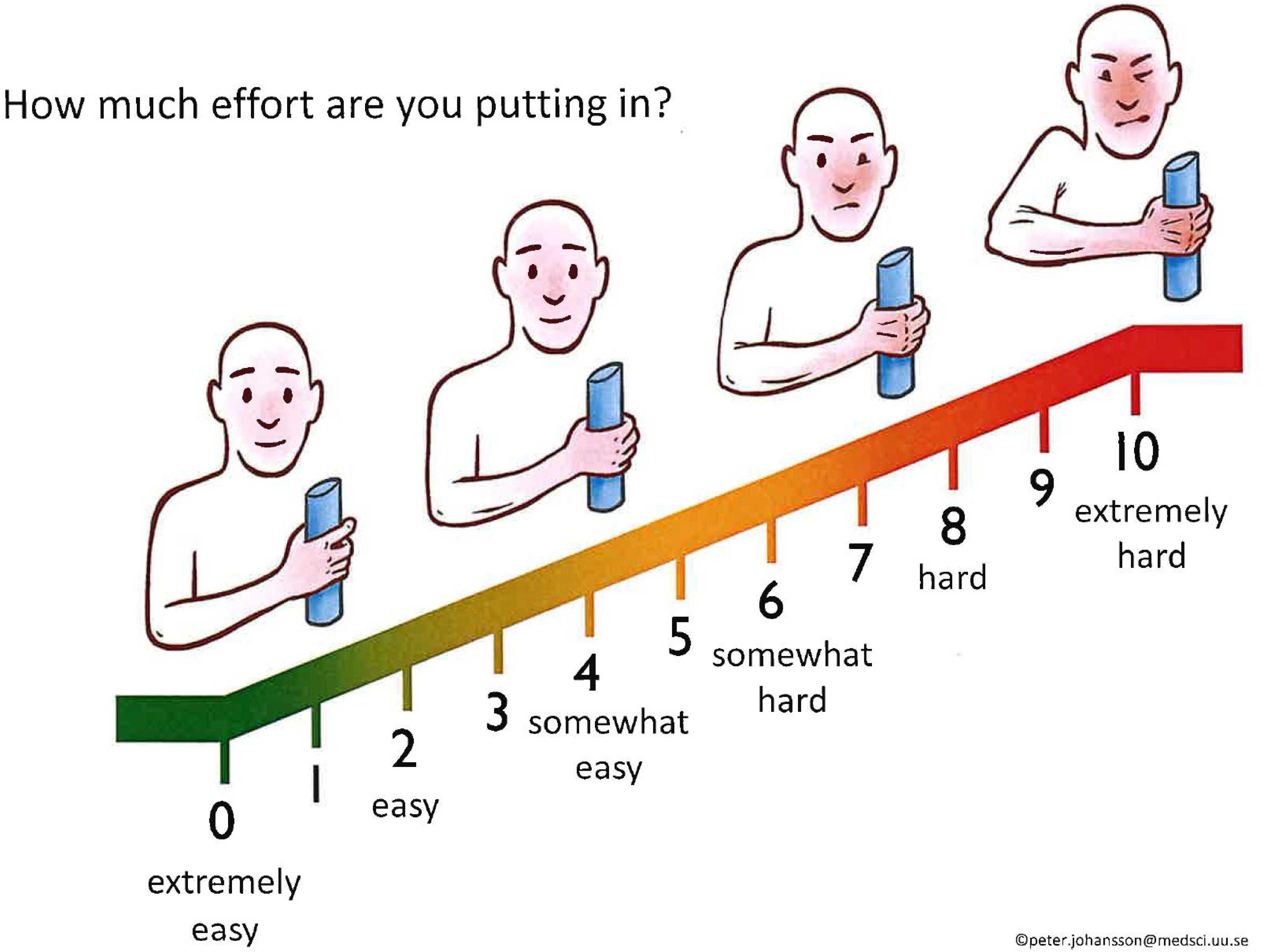
The modified OMNI-RES scale with the question, anchor words, and figures.

The detailed instructions for each task were described as follows:

1. For the **gripping task**, participants were seated in a chair, holding the dynamometer in their dominant hand, with their arm resting on the armrest and their elbow positioned at a 90-degree angle. The grip size of the dynamometer was adjusted to individual preferences for comfort.
2. For the **thumb-pinching task**, participants stood with their dominant arm hanging vertically and freely at their side. The pinch meter was placed on a table, which was customized to the appropriate height, allowing participants to press their thumb on the meter without the need to lean or raise their shoulders.
3. For the **screwing task**, participants stood holding a screwdriver in their dominant hand, with their elbow positioned at approximately a 100-degree angle. The screwdriver was inserted into a nozzle connected to the moment meter, which was securely fixated on a table adjusted to the participant’s height. During the exertion, participants turned the screwdriver outward, with the direction of the moment standardized to represent outward movement, regardless of which hand was used.
4. For the **key-pinching task**, participants were seated in a chair, holding the meter between their thumb and index finger, as if holding a key. The elbow was positioned at a 90-degree angle, kept close to the body, and supported by the chair’s armrest.

During both thumb- and key-pinching tasks, participants were specifically instructed to use only the force from their hand and forearm, avoiding leaning or any use of body weight.

### 2.3 Measurement of Exerted Force

The exerted force during the gripping task was measured using a dynamometer (G200, Biometrics Ltd, Newport, UK) with a sampling rate of 50 Hz. The exerted moment (the term was used interchangeably as force for the convenience in this study) during the screwing task was measured using a moment meter (Mecmesin static moment meter ST, Chauvin Arnoux Mätsystem, Täby, Sweden) at 100 Hz, with the direction of the moment standardized. Both the key-pinching and thumb-pinching forces were quantified using a pinch meter (P200, Biometrics Ltd, Newport, UK) at 100 Hz.

### 2.4 Measurement of Muscle activity

Muscle activity was recorded using self-adhesive bipolar electrodes with gel (Ag/AgCl electrodes, N-00-S/25, Ambu, Penang, Malaysia) and captured by a data logger (DataLog MWX8, Biometrics Ltd, Newport, UK). Signals were sampled at 1000 Hz per channel using a 24-bit A/D converter embedded in the logger.

Three sEMG placements were used: extensor, flexor, and through-forearm (Figure 3). The extensor sEMG was placed one-third of the forearm’s length from the elbow, above the area surrounding the m. extensor carpi radialis longus and brevis, with a center-to-center electrode distance of 2 cm (Figure 3a). The flexor sEMG was placed over the belly of the m. flexor digitorum superficialis, also with a center-to-center distance of 2 cm (Figure 3b) (Takala & Toivonen, 2013). For the through-forearm placement (Figure 3a,b), one of the bipolar electrodes was positioned over the m. extensor digitorum communis, while the other electrode was placed 1.5 cm lateral to the flexor sEMG electrodes (Takala & Toivonen, 2013).

**Figure 3.**
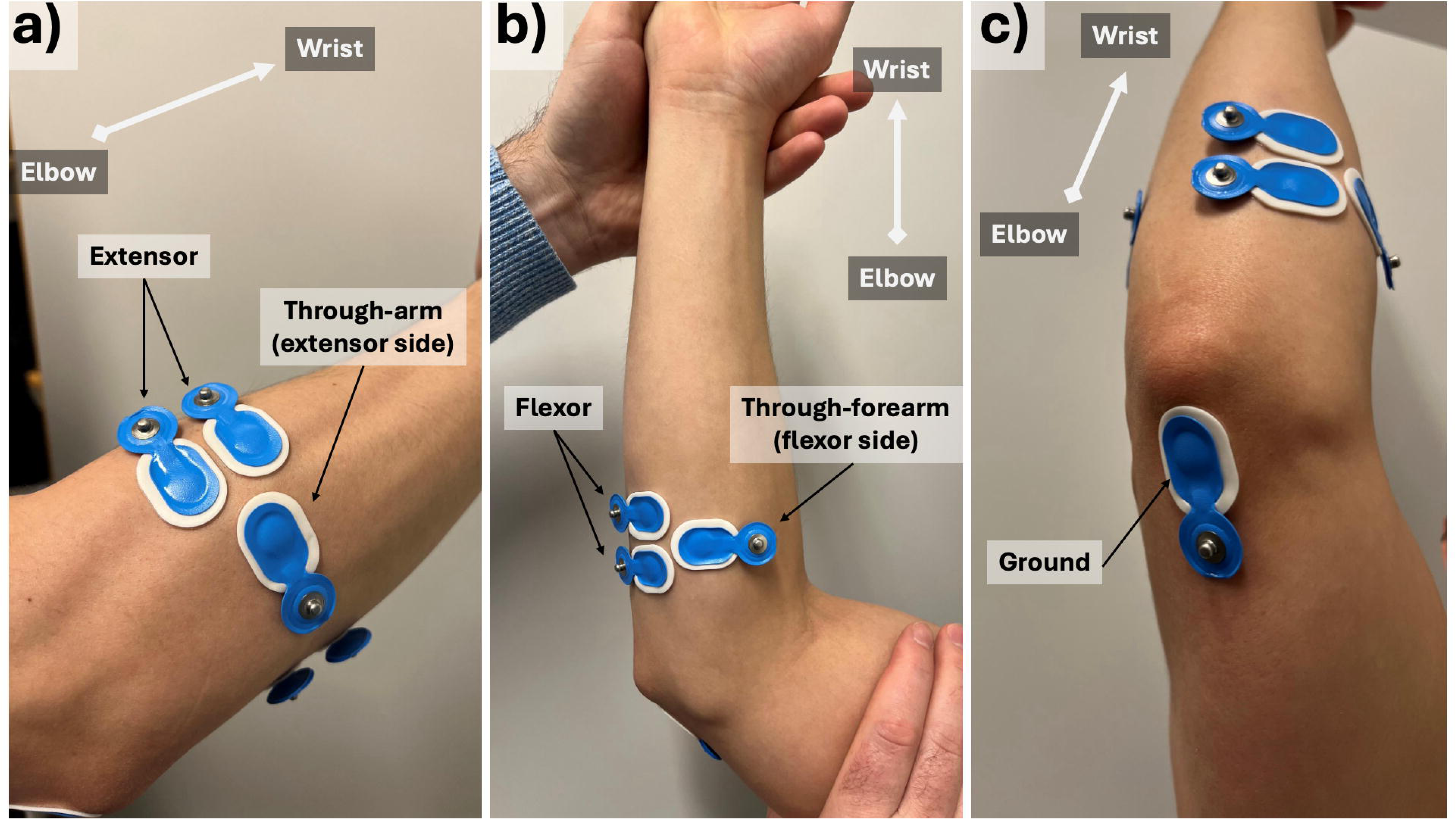
The placements of the sEMG electrodes on the right arm. Each placement is illustrated as such: extensor (a), flexor (b), through-forearm (a,b), and ground (c). The orientation of the arm is indicated by a light-grey arrow.

Ground electrodes were positioned above the olecranon process on both the distal forearm and humeral areas (Figure 3c).

### 2.5 Signal processing

The sEMG measurement was processed following a modified version of a previous study (Fan et al., 2022; Hansson et al., 2000). The sEMG signals were initially filtered using a digital bandpass filter (20–400 Hz) to isolate relevant frequencies, followed by a notch filter to remove environmental noise at 50 Hz and its harmonics. The resulting signals formed the base sEMG data. The noise level was determined as the minimum root mean square (RMS) value calculated from the base sEMG data during the resting period of the calibration phase, using a 2.375-second moving RMS window on the processed data. To determine each participant’s maximal voluntary electrical activity (MVE), the base sEMG data from the gripping task’s MVC period was extracted and filtered using a 0.5-second moving RMS window. The filtered data were then power-subtracted to remove noise using the formula 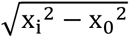, where x_0_ represents the noise level. The highest RMS value from the resulting data during the gripping MVC period was recorded as the participant’s MVE. To calculate muscle activity levels during each task and exertion level, the base sEMG data for each exertion level was first rectified using a 0.125-second moving RMS window, with the noise level similarly power-subtracted. The resulting sEMG data were then normalized to the individual’s MVE and expressed as a percentage of MVE (%MVE), representing the normalized muscle activity. The muscle activity level for each exertion level was obtained as the median value of the normalized muscle activity during the corresponding exertion period.

Similarly, force data were processed by an 8 Hz low-pass filter to the raw signals and saved as the base force data. The base force data were then denoised by subtracting the minimal value during the 10-second preparation period at the beginning of each task and normalized to the maximal force (MVC) observed during the corresponding MVC period of each task, expressed as a percentage of MVC (%MVC).

### 2.6 Statistics

All analyses related to muscle activity (%MVE), perceived exertion (OMNI-RES score), and exerted force (%MVC) were done within subgroups stratified by task.

The descriptive results of muscle activity were first presented, which include the group median and quartiles of muscle activity of each sEMG placement within each exertion level under each task. To test if there were significant differences of muscle activity measured by the three EMG placements under each perceived exertion level, Friedman tests were used. The core analyses in this study were two-fold:

1. Perceived exertion and exerted force were estimated using three sEMG placements. Following the approach of a previous study (Barański et al., 2024), polynomial mixed-effects models were employed to estimate both outcomes. In these models, muscle activity was treated as the fixed effect, while subjects were modeled as random effects. The assumption of homogeneity was evaluated using Levene’s test, which was satisfied. The Shapiro-Wilk test indicated violations of normality in approximately one-third of the fitted models. Still, mixed-effects models are considered generally robust to such violations (Schielzeth et al., 2020), which allows the analysis to proceed. Polynomial terms were added progressively to the models, and model comparisons were performed to determine the necessity of higher-order terms. The final models are structured as follows:

a. Perceived Exertion Model:

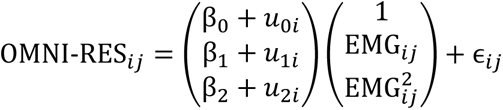

where OMNI-RES*_ij_* represents the perceived exertion for the subject *i* and task *j*, EMG*_ij_* is the muscle activity level, *β*_0_, *β*_1_, *β*_2_ are the fixed effects coefficients, and *u*_0*i*_, *u*_1*i*_, *u*_2*i*_ are the random effects for the subject *i*. The term *∈_ij_* denotes the residual error.
b. Exerted Force Model:

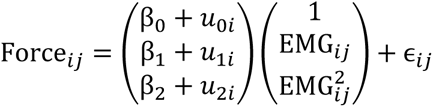

where Force*_ij_* represents the exerted force for the subject *i* and task *j*, EMG*_ij_* is the muscle activity level, *β*_0_, *β*_1_, *β*_2_ are the fixed effects coefficients, and *u*_0*i*_, *u*_1*i*_, *u*_2*i*_ are the random effects for the subject *i*. The term *∈_ij_* denotes the residual error. After fitting, the inter-task variances of the coefficients of the models were calculated to estimate the consistence of models from each sEMG placement across different tasks.
2. To explore the relationship between the three sEMG placements, Pearson correlations were calculated between each pair of sEMG placements, both within each task and across all tasks. Afterward, a secondary exploration was conducted by examining the contribution of extensor and flexor activities to through-forearm activity. To do so, a linear mixed-effects model was used, with z-score transformed muscle activity levels (calculated within each task of each participant):

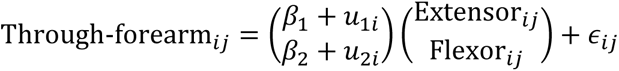

where Through-forearm*_ij_*, Extensor*_ij_*, Flexor*_ij_* represent the through-forearm, extensor, flexor muscle activity level for subject *i* and task *j*, *β*_1_, *β*_2_ are the fixed effects coefficients, and *u*_1*i*_, *u*_2*i*_ are the random effects for subject *i*. Intercepts were excluded under the assumption that when both muscles’ activities are zero, the through-forearm should be simultaneously zero. The term *∈_ij_* denotes the residual error.

## 3 Results

### 3.1 Prediction of Perceived Exertion from sEMG Placements

Overall, muscle activity from all three sEMG placements increased monotonically with rising perceived exertion levels. Among the placements, muscle activity from the extensor was consistently higher than that of the flexor across all tasks and perceived exertion levels. The through-forearm placement showed muscle activity that was intermediate between the extensor and flexor placements at lower perceived exertion levels across all four tasks. However, at higher perceived exertion levels, the through-forearm placement exhibited greater muscle activity than both the extensor and flexor placements during the screwing and thumb-pinching tasks, while remaining intermediate for the gripping and key-pinching tasks. (Table 2)

**Table 2.** Group medians of the 50th percentile muscle activity at different perceived exertion levels from the three sEMG placements across various tasks (N=16).

| Task | Perceived exertion level | Extensor (%MVE) |  | Flexor (%MVE) |  | Through-forearm (%MVE) |  | Friedman | Ext vs Fle | Ext vs Thr | Fle vs Thr |
| --- | --- | --- | --- | --- | --- | --- | --- | --- | --- | --- | --- |
|  |  | Median [Q1-Q3] |  | Median [Q1-Q3] |  | Median [Q1-Q3] |  | p | p | p | p |
| Grip | 0 (Rest) | 7.2 | [5.0–11.4] | 1.1 | [0.4–2.0] | 2.5 | [1.6–3.4] | <0.001 | <0.001 | 0.008 | 0.040 |
|  | 2 (Easy) | 12.9 | [7.0–19.4] | 3.1 | [1.8–6.3] | 5.7 | [3.2–7.4] | <0.001 | <0.001 | 0.024 | 0.024 |
|  | 4 (Moderate) | 29.0 | [15.0–31.5] | 11.5 | [8.5–13.6] | 14.5 | [9.8–17.1] | <0.001 | <0.001 | 0.102 | 0.102 |
|  | 8 (Hard) | 42.4 | [27.6–53.7] | 28.6 | [17.1–53.0] | 34.5 | [22.1–46.9] | 0.444 | 0.648 | 1.000 | 1.000 |
|  | 10 (Maximum) | 82.7 | [80.9–87.0] | 85.1 | [81.3–86.4] | 83.8 | [80.0–86.6] | 0.779 | 1.000 | 1.000 | 1.000 |
| Key-pinch | 0 (Rest) | 2.2 | [1.3–3.6] | 0.3 | [0.2–0.9] | 1.3 | [0.9–1.5] | <0.001 | <0.001 | 0.472 | 0.001 |
|  | 2 (Easy) | 6.6 | [3.7–12.8] | 0.7 | [0.5–2.0] | 3.1 | [1.9–3.9] | <0.001 | <0.001 | 0.040 | 0.008 |
|  | 4 (Moderate) | 14.2 | [7.4–20.2] | 2.3 | [1.5–7.2] | 7.3 | [5.1–11.6] | <0.001 | <0.001 | 0.335 | 0.004 |
|  | 8 (Hard) | 25.8 | [18.2–33.6] | 7.9 | [4.2–12.9] | 19.0 | [12.6–23.9] | <0.001 | <0.001 | 1.000 | 0.002 |
|  | 10 (Maximum) | 46.8 | [36.7–66.6] | 18.9 | [15.3–22.9] | 42.2 | [33.2–54.9] | <0.001 | <0.001 | 1.000 | <0.001 |
| Screw | 0 (Rest) | 4.6 | [2.7–6.8] | 1.9 | [1.3–3.1] | 2.7 | [2.1–6.0] | 0.002 | 0.001 | 0.231 | 0.231 |
|  | 2 (Easy) | 12.6 | [6.0–16.9] | 10.8 | [6.4–17.3] | 10.4 | [7.2–18.5] | 0.829 | 1.000 | 1.000 | 1.000 |
|  | 4 (Moderate) | 20.5 | [10.2–30.5] | 20.0 | [15.2–35.6] | 27.3 | [18.8–43.0] | 0.010 | 0.648 | 0.008 | 0.231 |
|  | 8 (Hard) | 36.0 | [28.4–56.9] | 49.0 | [35.4–63.7] | 59.4 | [44.9–78.3] | 0.010 | 0.231 | 0.008 | 0.648 |
|  | 10 (Maximum) | 74.0 | [60.9–91.0] | 77.5 | [60.8–92.2] | 93.3 | [85.5–107.4] | 0.050 | 1.000 | 0.102 | 0.102 |
| Thumb-pinch | 0 (Rest) | 1.2 | [0.4–2.4] | 0.5 | [0.3–0.7] | 1.2 | [1.0–1.5] | <0.001 | <0.001 | 1.000 | <0.001 |
|  | 2 (Easy) | 4.6 | [2.3–7.0] | 2.6 | [1.2–4.0] | 4.4 | [2.9–8.1] | 0.039 | 0.231 | 1.000 | 0.040 |
|  | 4 (Moderate) | 8.8 | [4.9–14.3] | 5.8 | [4.2–9.5] | 10.9 | [7.0–16.2] | 0.028 | 0.335 | 0.867 | 0.024 |
|  | 8 (Hard) | 19.5 | [16.3–26.9] | 16.9 | [11.6–22.3] | 26.4 | [18.5–33.5] | 0.039 | 1.000 | 0.231 | 0.040 |
|  | 10 (Maximum) | 47.0 | [31.1–72.2] | 32.8 | [25.5–42.3] | 57.2 | [43.7–62.3] | 0.099 | 0.231 | 1.000 | 0.155 |

When predicting perceived exertion levels from muscle activity, the polynomial mixed-effects models for all sEMG placements explained over 80% of the variance across all tasks. Notably, the model based on the through-forearm placement consistently explained more than 90% of the variance, with three tasks reaching around 95%, outperforming the other two placements. (Figure 4)

**Figure 4.**
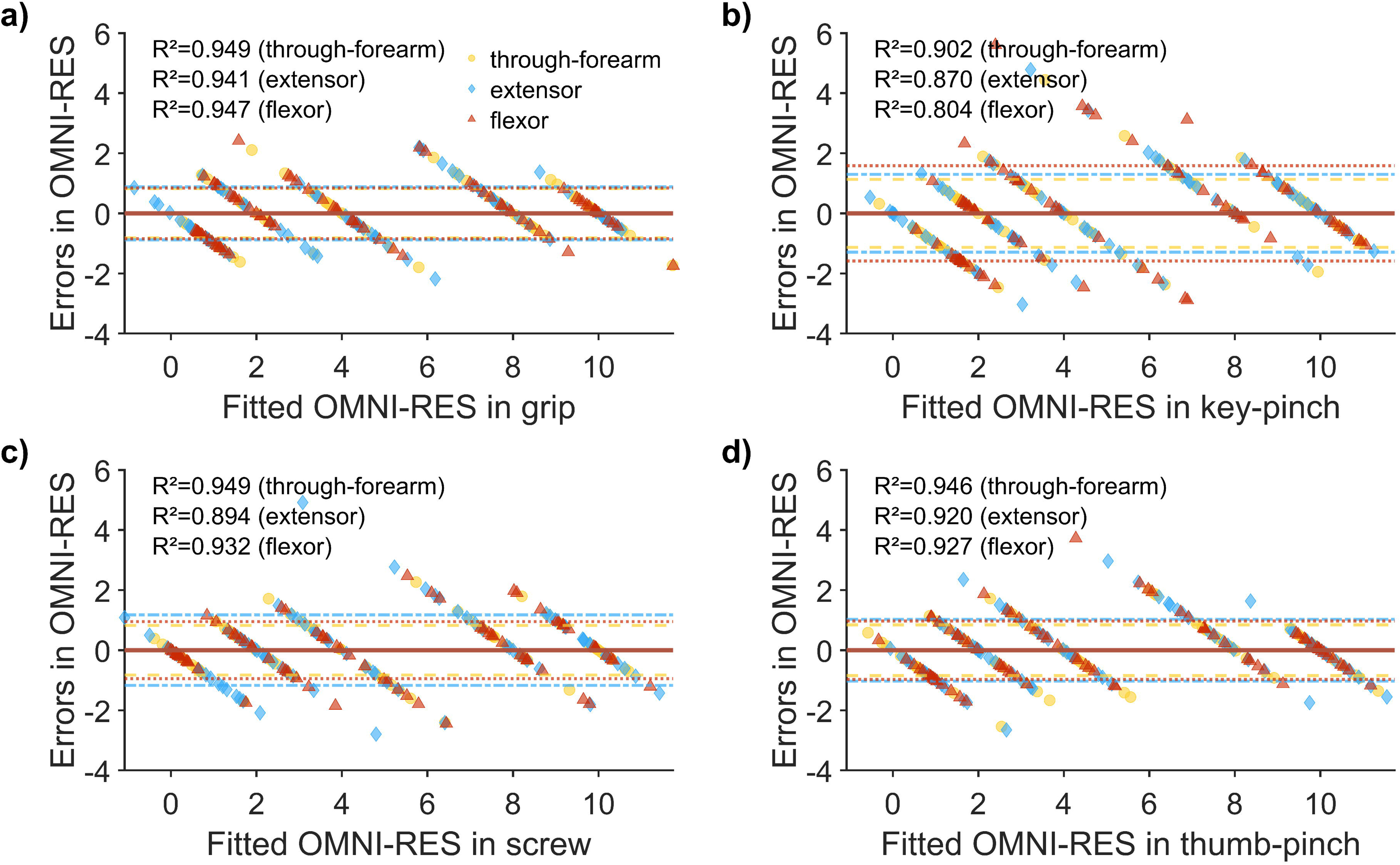
Residual plots of the mixed-effects models predicting perceived exertion levels (shown as OMNI-RES scores) from muscle activity across the four measured tasks.

The standard deviations (SDs) of the residuals were approximately 1 for the gripping, screwing, and thumb-pinching tasks, and less than 2 for the key-pinching task. When comparing the sEMG placements, the through-forearm placement exhibited the lowest residual SDs, indicating more precise predictions. (Figure 4)

To compare the consistency of models across different tasks and subjects, the inter-task variances of the model coefficients and the random factor coefficients were shown in Table S1 (Appendix). Overall, using the fixed effects as references, models from all three sEMG placements exhibited from low to moderate inter-task (1% – 66%) and inter-subject (22% – 75%) variability (excluding few extreme values for the insignificant intercepts). However, the differences between the placements were inconsistent. No single sEMG placement consistently outperformed the others; each showed strengths in certain tasks but underperformed in others.

### 3.2 Predicting force exertion from sEMG placements

The prediction of exerted force from muscle activity showed strong fits of models across all four tasks, with explained variances (R^2^) mostly exceeding 95% (0.95) and most SDs of the residuals less than 10 %MVC. Among the three sEMG placements, models based on the through-forearm sEMG consistently explained the largest proportion of variance across all tasks, with three tasks achieving R^2^>0.98 and the key-pinching task reaching R^2^>0.96, albeit marginally. Additionally, model residuals from the through-forearm placement were the lowest across all tasks, with three tasks showing residuals below 5 %MVC and the key-pinching task below 8 %MVC. (Figure 5)

**Figure 5.**
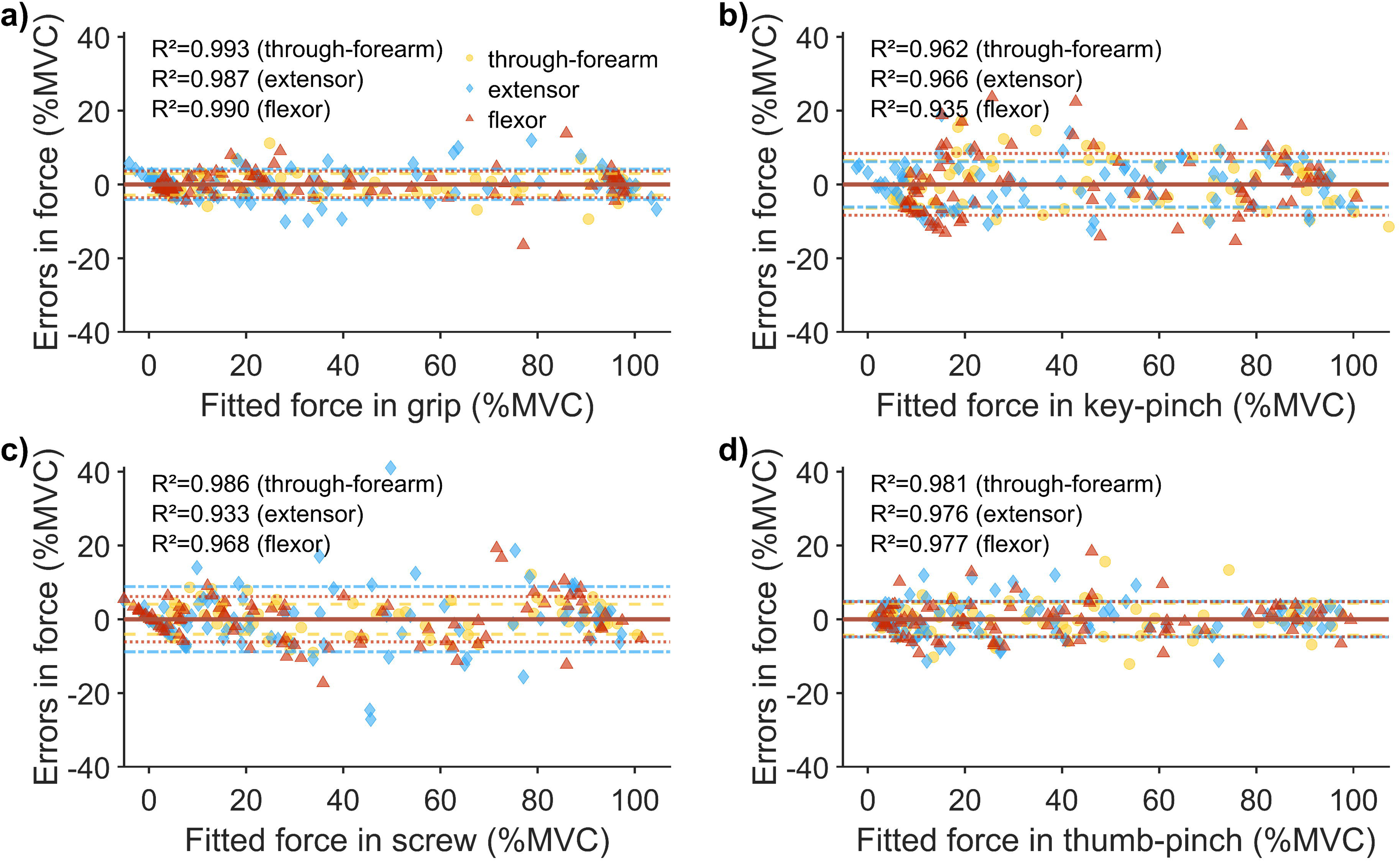
Residual plots of the mixed-effects models predicting exerted force from muscle activity across the four measured tasks.

Despite the high R^2^ values and low residuals observed across individual models, inter-subject and inter-task variability was prominent in all three sEMG placements. For instance, in the model for the extensor during gripping, the SD of the slope due to the random effect (subject) was 0.87, which accounted for 48% of the fixed-effect slope (1.82), indicating substantial variability between individuals. Similarly, task-specific differences were evident, as seen in the extensor model, where the slope during key-pinching (3.83) was more than double that during gripping. (Table S2 in Appendix)

Despite the noticeable variability, among the three sEMG placements, the through-forearm placement demonstrated relatively low inter-subject variability (20% – 73% of the corresponding fixed effects) across most tasks, and in some cases, it exhibited the lowest variability, outperforming the extensor (18% – 103%) and flexor (26% – 152%) placement. Additionally, the through-forearm (2%–91%) and extensor (2%–98%) placement showed a similarly low to moderate inter-task variability ratio relative to gripping for the fixed factors, while the flexor placement had the highest ratio in all tasks (137%–626%). (Table S2 in Appendix)

### 3.3 Correlations between sEMG placements

Further analysis of the correlations between the 50th percentile muscle activity revealed strong linear relationships across all three sEMG placements in all four tasks (all r>0.97). (Table 3)

**Table 3.**
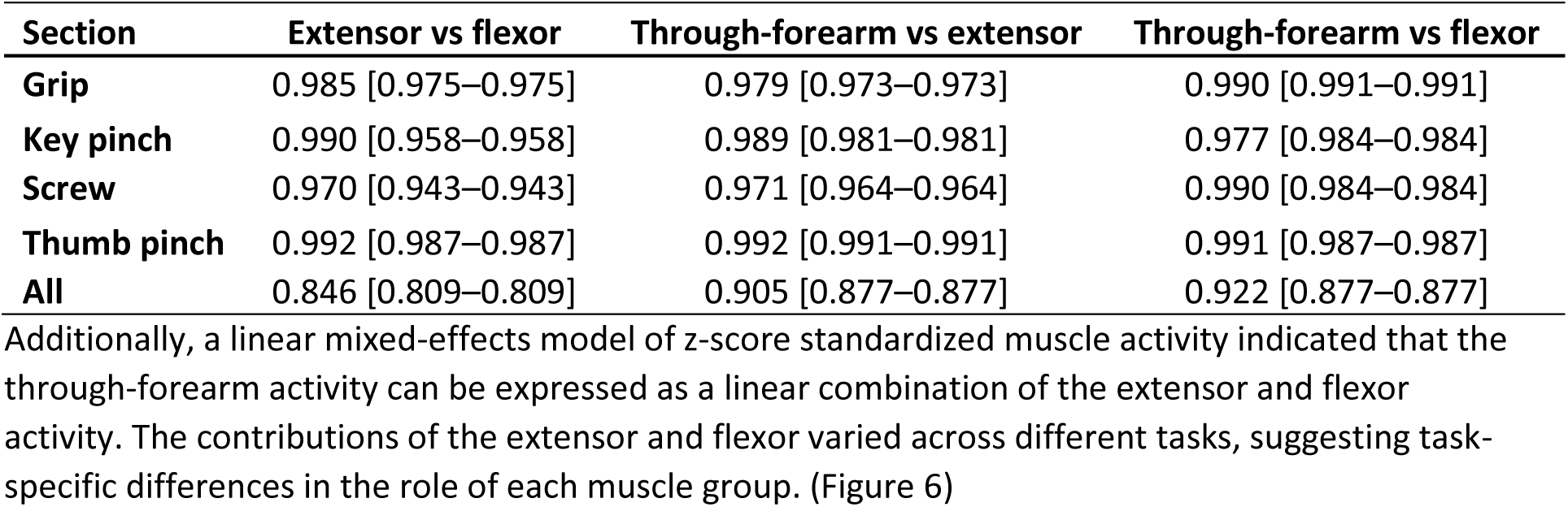
Pearson correlations of the 50^th^ percentile muscle activity between the three sEMG placements across different tasks.

| Section | Extensor vs flexor | Through-forearm vs extensor | Through-forearm vs flexor |
| --- | --- | --- | --- |
| **Grip** | 0.985 [0.975–0.975] | 0.979 [0.973–0.973] | 0.990 [0.991–0.991] |
| **Key pinch** | 0.990 [0.958–0.958] | 0.989 [0.981–0.981] | 0.977 [0.984–0.984] |
| **Screw** | 0.970 [0.943–0.943] | 0.971 [0.964–0.964] | 0.990 [0.984–0.984] |
| **Thumb pinch** | 0.992 [0.987–0.987] | 0.992 [0.991–0.991] | 0.991 [0.987–0.987] |
| **All** | 0.846 [0.809–0.809] | 0.905 [0.877–0.877] | 0.922 [0.877–0.877] |

Additionally, a linear mixed-effects model of z-score standardized muscle activity indicated that the through-forearm activity can be expressed as a linear combination of the extensor and flexor activity. The contributions of the extensor and flexor varied across different tasks, suggesting task-specific differences in the role of each muscle group. (Figure 6)

**Figure 6.**
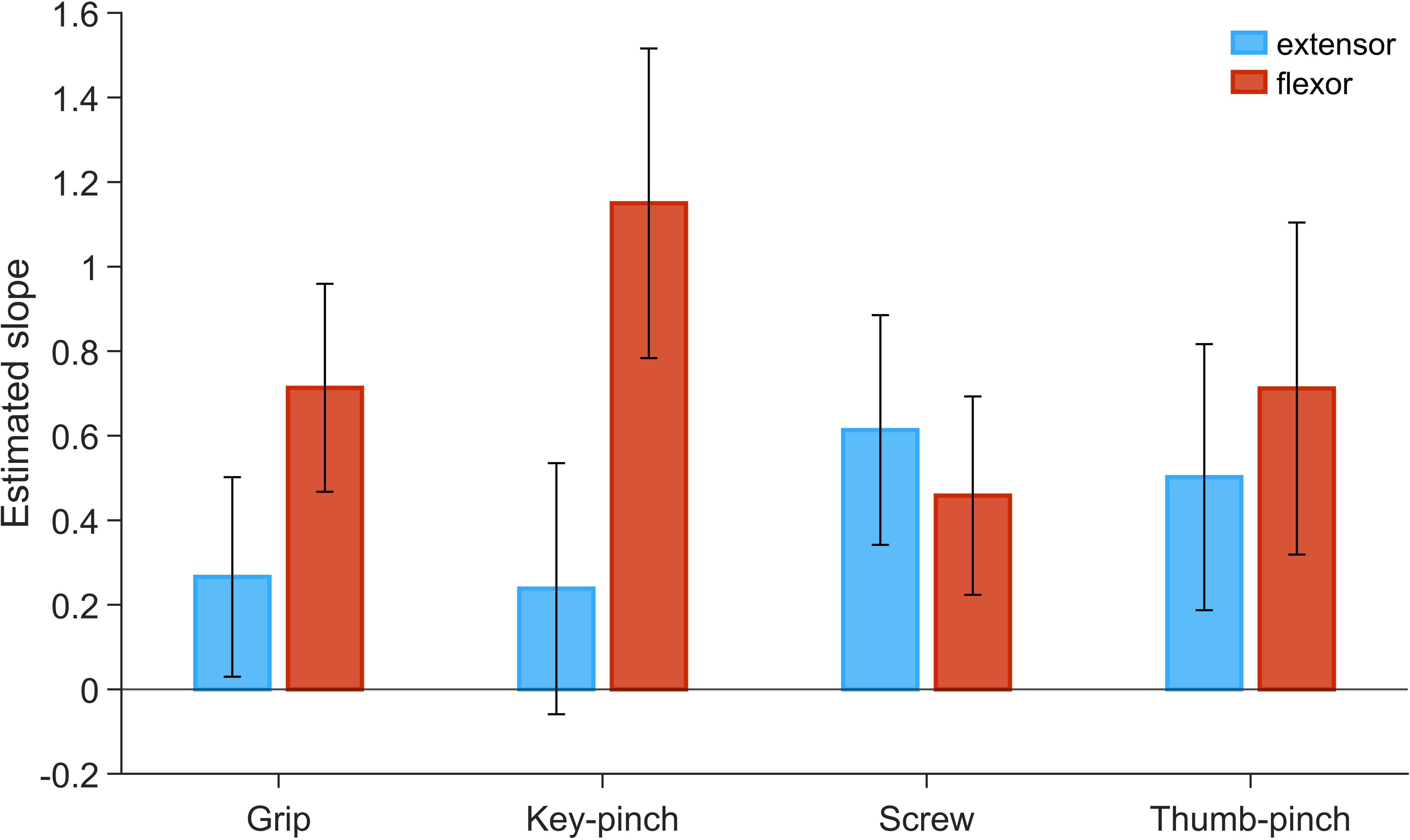
Slopes of the extensor and flexor activity in the z-score standardized linear mixed-effects models for the through-forearm activity across different tasks. Whiskers represent the lower and upper boundaries of the estimated slopes.

## 4 Discussion

This study highlights that sEMG can effectively predict both perceived exertion (i.e., OMNI-RES levels) and normalized exerted force with good fitness and low errors. Among the three sEMG placements evaluated, the through-forearm placement consistently explained the largest proportion of variance and produced the lowest residuals across all four tasks, making it a reliable option for estimating both perceived exertion and exerted force. Despite notable inter-subject and inter-task variability observed across all placements, the through-forearm placement demonstrated the lowest inter-subject variability in most tasks, outperforming the extensor and flexor placements in this regard.

Additionally, strong correlations were observed among the three sEMG placements—extensor, flexor, and through-forearm. Specifically, through-forearm muscle activity was closely correlated with a linear combination of extensor and flexor activity, with the relative contributions of each varying depending on the task.

### 4.1 Predicting Perceived Exertion and Exerted Force

Our findings demonstrate that through-forearm sEMG can predict perceived exertion and normalized exerted force with high fitness (R^2^ > 96%) and low errors (<5% MVC for three tasks, and <8% MVC for key-pinching) across all placements and tasks, showcasing its utility of monitoring exertions in work tasks.

Similar results have been observed in previous studies. Barański et al. reported that when predicting grip force using a single sEMG channel placed on the flexor, models based on RMS yielded high R^2^ values (>0.8), with prediction errors below 10N across a 20–100N force range (Barański et al., 2024). Although their study focused on absolute force rather than %MVC, the results align closely with ours.

However, inter-task variability remains a critical factor, as evidenced by the relative variance in model parameters across tasks, particularly when comparing tasks like gripping and key-pinching. In practical settings, tasks often involve complex transitions between sub-tasks, which may reduce prediction accuracy. While the current data and results cannot precisely determine the extent to which task order affected the outcomes, the significant differences in parameter values between tasks—and the fact that these differences did not follow a monotonic trend along the temporal sequence—strongly suggest the presence of inter-task variability.

Several studies have attempted to mitigate this by increasing the number of sEMG channels to capture broader muscle activity patterns. For example, Wang et al. (2023) employed five sEMG channels to identify the task and predict exerted force, achieving a 74% precision in force category prediction. Bardizbanian et al. (2020) used a 12-channel sEMG armband, yielding an error rate of 2.5% to 8% MVC during finger contractions. Similarly, Chihara and Sakamoto (2021) utilized an 8-channel armband device, estimating container weights (ranging from 2 to 20 kg) with average absolute errors of 1.5 kg. However, Martinez et al. (2020) demonstrated that increasing the channel density beyond 16 may not significantly improve prediction accuracy. For applications where fewer channels are desired, alternative methods, such as integrating sensors like accelerometers (Mao, Zheng, et al., 2023) or pre-defining primary task types, may offer solutions. In clinical studies, the simplicity and cost-effectiveness of sensors often determine the feasibility of including more participants, further emphasizing the need for efficient methods in predicting force during real tasks.

Besides inter-task variability, inter- and intra-subject variability in the prediction of perceived exertion and exerted force is also important to consider. Although normalization has partially reduced this variability, individual differences in muscle engagement during the same tasks remain. Studies by Rezende et al. (2023) and Jasque-Bou et al. (2024) have shown that muscle activity patterns can vary significantly inter- and intra-subject even under identical force exertion and task conditions. These findings highlight the inevitable presence of both inter- and intra-subject variability. While normalization by MVCs helps, it may not suffice entirely. Further research is needed to explore alternative normalization techniques or methods to minimize the impact of this variability on research outcomes.

### 4.2 Interrelation Between sEMG Placements

Further analysis revealed that through-forearm activity was linearly correlated with both extensor and flexor activity. This finding is consistent with previous studies on muscle synergy in humans, where signals from the central nervous system are distributed across muscles according to a pre-learned synergy pattern (Bizzi & Cheung, 2013; Geng et al., 2020). According to the theory, different muscles were activated in a predetermined combination in various tasks.

In our study, through-forearm placement appears to capture a fraction of these synergies, allowing it to predict perceived exertion and exerted force slightly better than the other placements. Takala’s previous study also suggested a similar trend, with through-forearm placement exhibiting better correlation for certain tasks.

### 4.3 Methodological consideration

Compared to Takala’s work (Takala & Toivonen, 2013), our study expanded the range of tasks, allowing for greater generalization of results to common work-related activities. Fatigue is known to impact force prediction (J. Wang et al., 2021). While the order of tasks in our study was not randomized, several measures were taken to mitigate potential fatigue effects. First, participants rested for several minutes between tasks, providing sufficient time for muscle recovery and limiting the influence of fatigue on results. Second, the fixed task order alternated between pinching movements, which primarily engage hand muscles, and gripping and screwing tasks, which emphasize forearm muscles, allowing specific muscle groups to rest between exertions. Third, comparisons between sEMG placements were performed within each task, ensuring consistent pre-task workloads and controlling for fatigue-related variability.

The quadratic models selected through an incremental model comparison process produced promising results, aligning with previous research (e.g., Barański et al. (2024)). This incremental approach effectively balanced the risks of overfitting with the need to capture the non-linear relationship between perceived exertion and muscle activity. Our study also benefits from a demographically balanced dataset, which minimizes potential biases induced by demographic factors.

Additionally, our findings reveal that the through-forearm placement reflects a task-dependent, linear combination of extensor and flexor activity. This provides valuable insight into the nature of through-forearm measurements, suggesting that they capture integrated muscle activity across multiple muscle groups while varying according to task demands.

### 4.4 Limitations

One limitation of our study is that force was normalized by task, which may, therefore, influence the practicality of the resulting models. Although this approach helped control for task-specific variations, its impact on the predictive power of the models in real-world cases remains unclear. Furthermore, the number of tasks in this study was limited, and real-world hand movements are far more complex and diverse. In practice, hand movements often involve quick transitions between different motions, which may not be fully captured by our protocol. Even though our study aimed to expand the scope beyond simple gripping tasks, future studies will need to employ more innovative methods to capture the full dynamic range of hand movements.

### 4.5 Practical Implications

Our findings suggest that through-forearm sEMG can feasibly predict perceived and exerted force in controlled laboratory settings, consistently explaining the largest proportion of variance and producing the lowest residuals in predicting both perceived exertion and exerted force across all four tasks. However, strategies to account for inter-task and inter-subject variability are crucial. Acquiring task types for specific occupations at any stage of the measurement or conducting task-specific model calibrations could help mitigate inter-task variability. Regarding inter-subject variability, calibrations based solely on resting and maximal contractions may be insufficient. Incorporating normalization techniques, such as using progressively increasing contraction scales instead of a single maximal voluntary contraction, could enhance model accuracy.

Finally, if a single-channel sEMG approach is preferred, the choice of sEMG placement should be guided by the study’s objectives. For assessing muscle-specific workload, the sEMG should be placed on the target muscle (e.g., extensor or flexor). Whereas if the goal is to estimate overall workload or perceived exertion, the through-forearm placement may be more appropriate, as it captures signals from multiple muscles.

## 5 Conclusion

Our findings underscore the reliability of sEMG, particularly through-forearm placement, in estimating both perceived exertion and exerted force. However, limitations such as inter-task and inter-subject variability were identified, suggesting the need for further studies to explore compensatory methods for applying quadratic sEMG modeling on single-channel sEMG measurements in real-world settings. Additionally, the study suggests that through-forearm placement captures integrated muscle activity from multiple muscles, making it a slightly better option than muscle-specific placements for estimating physical workload in the hand and forearm. In practical applications, the choice of sEMG placement should align with the study’s specific objectives: through-forearm placement is better suited for general exertion predictions, while muscle-specific placements are more appropriate for assessing muscle-specific risks.

## 7 Appendix

**Table S1.** Variables and coefficients of the sEMG-perceived-exertion prediction models. Bold values indicate the fixed factors significantly improved the performance of the model.

| Task | Parameter | Extensor | Flexor | Through-forearm |
| --- | --- | --- | --- | --- |
| Grip | Fixed (Estimate): b0 | **-1.69** | **0.52** | -0.01 |
|  | Fixed (Estimate): sEMG | **0.33** | **0.34** | **0.33** |
|  | Fixed (Estimate): sEMG<sup>2</sup> | **-0.00229** | **-0.00265** | **-0.00250** |
|  | Random (SD): b0 | 0.28 | 0.07 | 0.13 |
|  | Random (SD): sEMG | 0.14 | 0.14 | 0.15 |
|  | Random (SD): sEMG<sup>2</sup> | 0.0017 | 0.0017 | 0.0017 |
|  | Residuals (SD) | 0.99 | 0.94 | 0.92 |
|  | R<sup>2</sup> (adjusted) | 0.941 | 0.947 | 0.949 |
| Screw | Fixed (Estimate): b0 | -0.43 | -0.27 | -0.16 |
|  | Fixed (Estimate): sEMG | **0.26** | **0.23** | **0.18** |
|  | Fixed (Estimate): sEMG<sup>2</sup> | **-0.00172** | **-0.00129** | **-0.00078** |
|  | Random (SD): b0 | 0.98 | 0.41 | 0.27 |
|  | Random (SD): sEMG | 0.05 | 0.09 | 0.06 |
|  | Random (SD): sEMG<sup>2</sup> | 0.0007 | 0.0009 | 0.0006 |
|  | Residuals (SD) | 1.31 | 1.06 | 0.92 |
|  | R<sup>2</sup> (adjusted) | 0.894 | 0.932 | 0.949 |
| Key-pinch | Fixed (Estimate): b0 | -0.15 | **1.16** | 0.41 |
|  | Fixed (Estimate): sEMG | **0.39** | **0.89** | **0.49** |
|  | Fixed (Estimate): sEMG<sup>2</sup> | **-0.00364** | **-0.0226** | **-0.00596** |
|  | Random (SD): b0 | 0.66 | 0.30 | 0.52 |
|  | Random (SD): sEMG | 0.10 | 0.24 | 0.11 |
|  | Random (SD): sEMG<sup>2</sup> | 0.0020 | 0.0085 | 0.0023 |
|  | Residuals (SD) | 1.44 | 1.73 | 1.27 |
|  | R<sup>2</sup> (adjusted) | 0.870 | 0.804 | 0.902 |
| Thumb-pinch | Fixed (Estimate): b0 | 0.37 | **0.56** | -0.03 |
|  | Fixed (Estimate): sEMG | **0.44** | **0.57** | **0.40** |
|  | Fixed (Estimate): sEMG<sup>2</sup> | **-0.00435** | **-0.00829** | **-0.00394** |
|  | Random (SD): b0 | 0.67 | 0.45 | 0.47 |
|  | Random (SD): sEMG | 0.16 | 0.17 | 0.09 |
|  | Random (SD): sEMG<sup>2</sup> | 0.0024 | 0.0039 | 0.0016 |
|  | Residuals (SD) | 1.15 | 1.12 | 0.95 |
|  | R<sup>2</sup> (adjusted) | 0.920 | 0.927 | 0.946 |
| Inter-task variances | Fixed (Estimate): b0 | 0.77 | 0.34 | 0.06 |
|  | Fixed (Estimate): sEMG | 0.01 | 0.09 | 0.02 |
|  | Fixed (Estimate): sEMG<sup>2</sup> | <0.0001 | <0.0001 | <0.0001 |
|  | Random (SD): b0 | 0.08 | 0.03 | 0.03 |
|  | Random (SD): sEMG | <0.01 | <0.01 | <0.01 |
|  | Random (SD): sEMG<sup>2</sup> | <0.0001 | <0.0001 | <0.0001 |

**Table S2.** Parameters and coefficients of the sEMG-Force prediction models. Bold values indicate the fixed factors significantly improved the performance of the model.

| Task | Parameter | Extensor | Flexor | Through-forearm |
| --- | --- | --- | --- | --- |
| **Grip** | Fixed (Estimate): b0 | **-11.06** | 1.28 | **-1.75** |
|  | Fixed (Estimate): sEMG | **1.82** | **1.99** | **1.88** |
|  | Fixed (Estimate): sEMG<sup>2</sup> | **-0.00654** | **-0.0104** | **-0.00858** |
|  | Random (SD): b0 | 4.72 | 0.16 | 0.58 |
|  | Random (SD): sEMG | 0.87 | 0.51 | 0.41 |
|  | Random (SD): sEMG<sup>2</sup> | 0.0099 | 0.0058 | 0.0048 |
|  | Residuals (SD) | 4.86 | 4.01 | 3.28 |
|  | R<sup>2</sup> (adjusted) | 0.987 | 0.990 | 0.993 |
| **Screw** | Fixed (Estimate): b0 | **-6.40** | -3.24 | -1.69 |
|  | Fixed (Estimate): sEMG | **1.96** | **1.45** | **1.06** |
|  | Fixed (Estimate): sEMG<sup>2</sup> | **-0.00917** | -0.00321 | -0.00106 |
|  | Random (SD): b0 | 5.81 | 4.34 | 1.96 |
|  | Random (SD): sEMG | 0.36 | 0.88 | 0.49 |
|  | Random (SD): sEMG<sup>2</sup> | 0.0027 | 0.0082 | 0.0042 |
|  | Residuals (SD) | 9.90 | 7.33 | 4.93 |
|  | R<sup>2</sup> (adjusted) | 0.933 | 0.968 | 0.986 |
| **Key-pinch** | Fixed (Estimate): b0 | **-3.96** | **6.85** | **3.36** |
|  | Fixed (Estimate): sEMG | **3.83** | **9.19** | **4.14** |
|  | Fixed (Estimate): sEMG<sup>2</sup> | **-0.0390** | **-0.257** | **-0.0451** |
|  | Random (SD): b0 | 1.07 | 3.37 | 0.33 |
|  | Random (SD): sEMG | 1.24 | 4.14 | 0.82 |
|  | Random (SD): sEMG<sup>2</sup> | 0.0228 | 0.2061 | 0.0136 |
|  | Residuals (SD) | 7.17 | 9.94 | 7.15 |
|  | R<sup>2</sup> (adjusted) | 0.966 | 0.935 | 0.962 |
| **Thumb-pinch** | Fixed (Estimate): b0 | 0.97 | **3.12** | -0.41 |
|  | Fixed (Estimate): sEMG | **3.52** | **4.17** | **2.65** |
|  | Fixed (Estimate): sEMG<sup>2</sup> | **-0.0306** | **-0.0595** | **-0.0187** |
|  | Random (SD): b0 | 1.21 | 1.49 | 1.48 |
|  | Random (SD): sEMG | 2.52 | 2.88 | 1.05 |
|  | Random (SD): sEMG<sup>2</sup> | 0.0315 | 0.0903 | 0.0137 |
|  | Residuals (SD) | 5.70 | 5.90 | 5.15 |
|  | R<sup>2</sup> (adjusted) | 0.976 | 0.977 | 0.981 |
| **Inter-task variances** | Fixed (Estimate): b0 | 25.12 | 17.58 | 5.77 |
|  | Fixed (Estimate): sEMG | 1.08 | 12.45 | 1.72 |
|  | Fixed (Estimate): sEMG<sup>2</sup> | 0.0003 | 0.0142 | 0.0004 |
|  | Random (SD): b0 | 5.87 | 3.51 | 0.58 |
|  | Random (SD): sEMG | 0.85 | 2.93 | 0.09 |
|  | Random (SD): sEMG<sup>2</sup> | 0.0002 | 0.0089 | <0.0001 |

